## Supplemental files for "Endocardial HDAC3 is required for myocardial trabeculation"

### Supplemental Materials

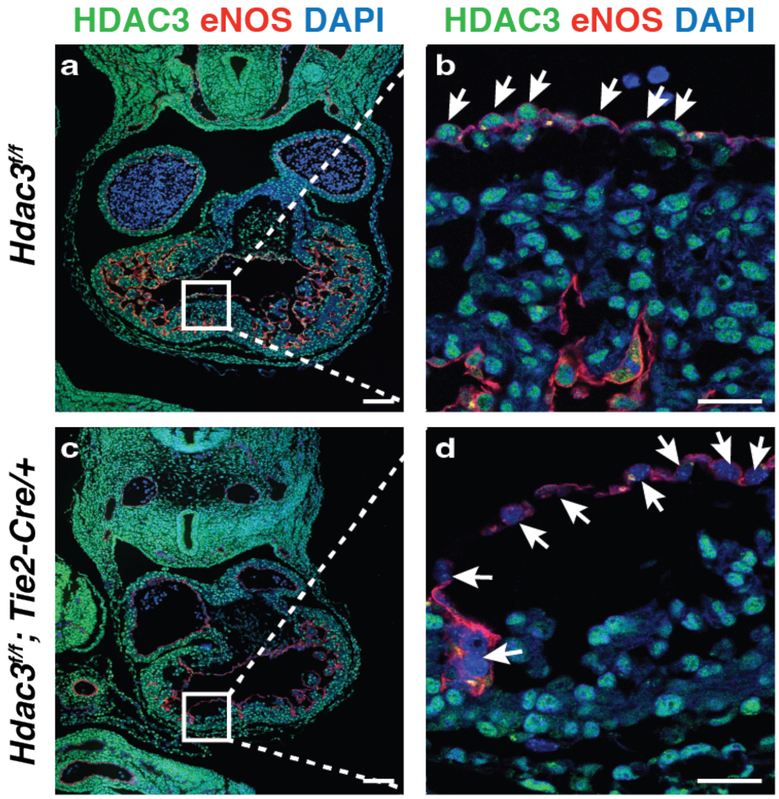


**Supplemental Figure 1. Restrictive ablation of *Hdac3* in the developing cardiac endothelial cells.**

Immunofluorescence staining shows specific deletion of HDAC3 in the cardiac endothelial cells (eNOS+, arrows) in E10.5 *Hdac3*$^{tko}$ (*Hdac3*$^{f/f}$; *Tie2-Cre/*+) heart as compared to the littermate control (*Hdac3*$^{f/f}$) heart (a). Scale bars, a and c, 100 μm; b and d, 25 μm.

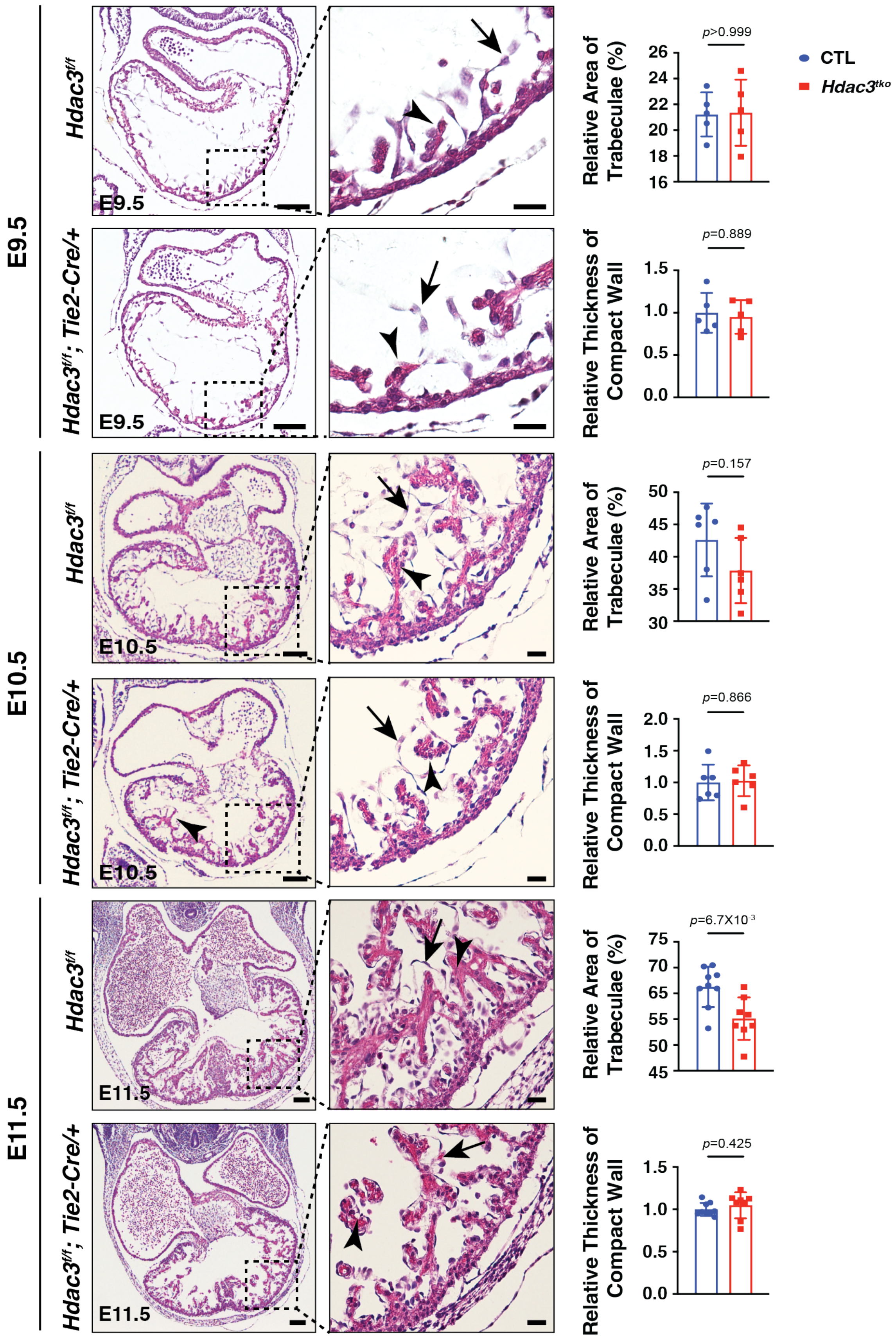
E9.5
E10.5
E11.5
Hdac3f/f
Hdac3f/f; Tie2-Cre/+
E9.5
E10.5
E11.5
Relative Area of Trabeculae (%)
Relative Thickness of Compact Wall
p>0.999
p=0.889
p=0.157
p=0.866
p=6.7X10-3
p=0.425
CTL
Hdac3tko

**Supplemental Figure 2. Histological analyses of cardiac phenotypes of *Hdac3*$^{tko}$ hearts at early embryonic stages.**

Hematoxylin and eosin staining on cross sections of E9.5-E11.5 *Hdac3*$^{tko}$ embryos (*Hdac3*$^{f/f}$*; Tie2-Cre/*+) and littermate control (CTL) embryos (*Hdac3*$^{f/f}$). Arrows mark endocardium and arrowheads point to trabeculae. Quantifications are shown on the right. For E9.5, CTL: n=5, *Hdac3*$^{tko}$: n=5; For E10.5, CTL: n=6, *Hdac3*$^{tko}$: n=6; For E11.5, CTL: n=9, *Hdac3*$^{tko}$: n=8. Scale bars, 100 μm and 25 μm (in insets). *P*-values were determined by the Mann-Whitney U test for E9.5 embryos, and unpaired two tailed Student's *t*-test for E10.5 and E11.5 embryos.

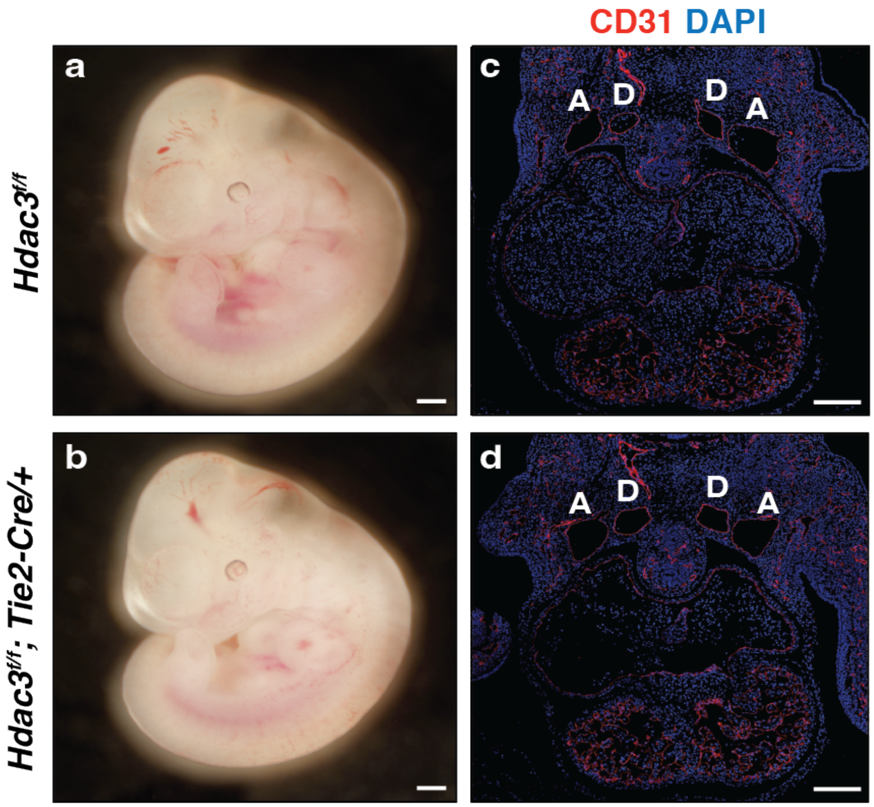


**Supplemental Figure 3. Normal vasculature in *Hdac3*$^{tko}$ embryos.**

a and b, gross morphology of an E11.5 *Hdac3*$^{tko}$ (*Hdac3*$^{f/f}$*; Tie2-Cre/*+) embryo (b) and an E11.5 littermate control (CTL, *Hdac3*$^{f/f}$) embryo (a). b and d, immunofluorescence staining of CD31. There were no apparent differences in either gross appearance or vasculature between *Hdac3*$^{tko}$ and CTL embryos. A, anterior cardinal vein; D, dorsal aorta. Scale bars, a&b, 500um; c&d, 200um.

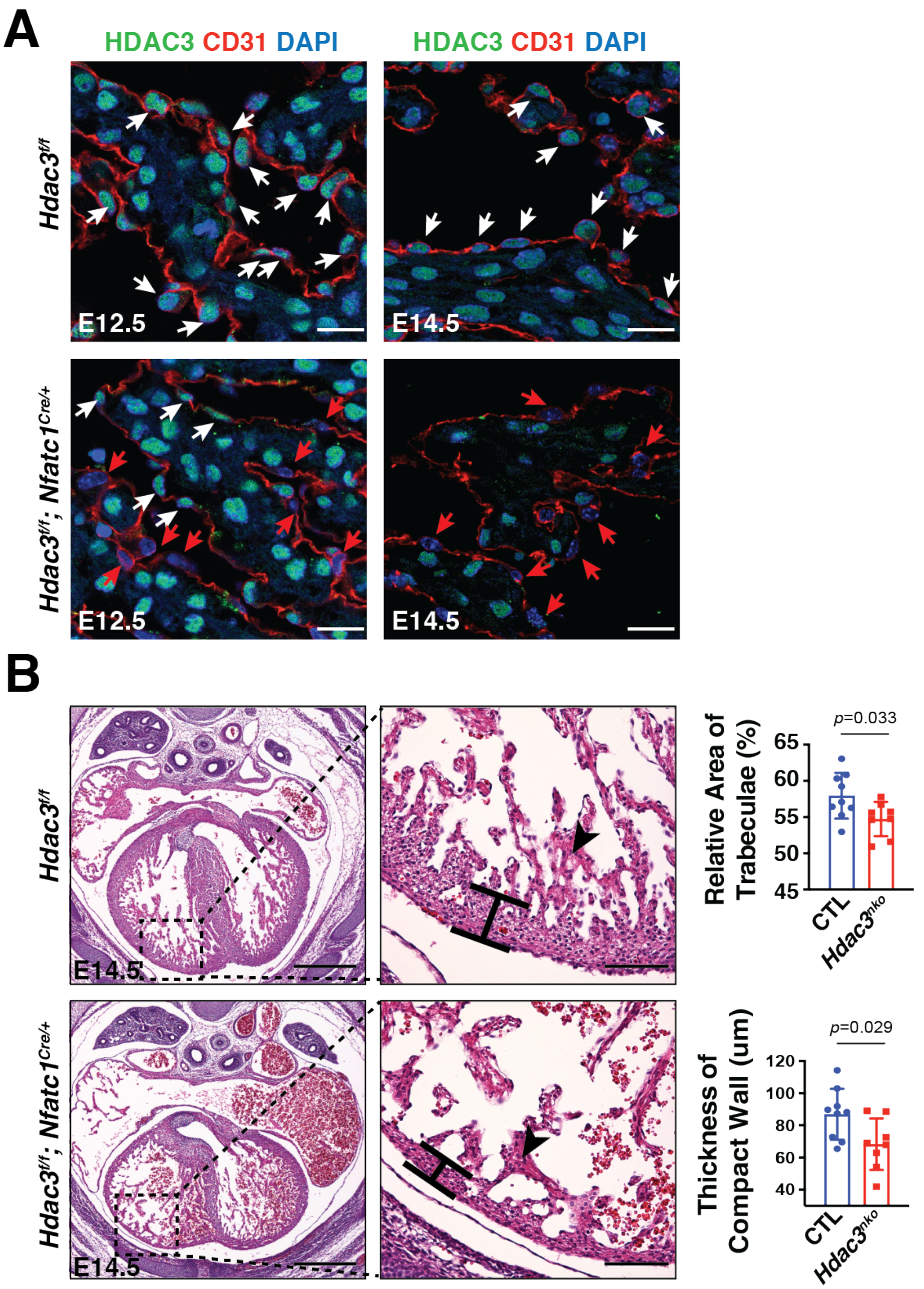


**Supplemental Figure 4. Endocardial specific deletion of *Hdac3* results in ventricular hypoplasia.**

**(A)** Deletion of *Hdac3* by *Nfatc1*$^{Cre/+}$ in the developing endocardium (E12.5 and E14.5). Immunofluorescence staining for HDAC3 and eNOS was performed. White arrows point to endocardial cells that express HDAC3, whereas red arrows point to endocardial cells in which HDAC3 is absent. Scale bars, 20 μm. **(B)** Cardiac phenotypes of E14.5 *Hdac3* endocardial knockout (*Hdac3*$^{nko}$, *Hdac3*$^{f/f}$*; Nfatc1*$^{Cre/+}$) embryos. Arrowheads point to trabeculae. Scale bars, 500 μm (main panels) and 100 μm (insets). Quantifications are shown on the right. CTL: n=9, *Hdac3*$^{nko}$: n=8. *P*-values were determined by unpaired two tailed Student's *t*-test.

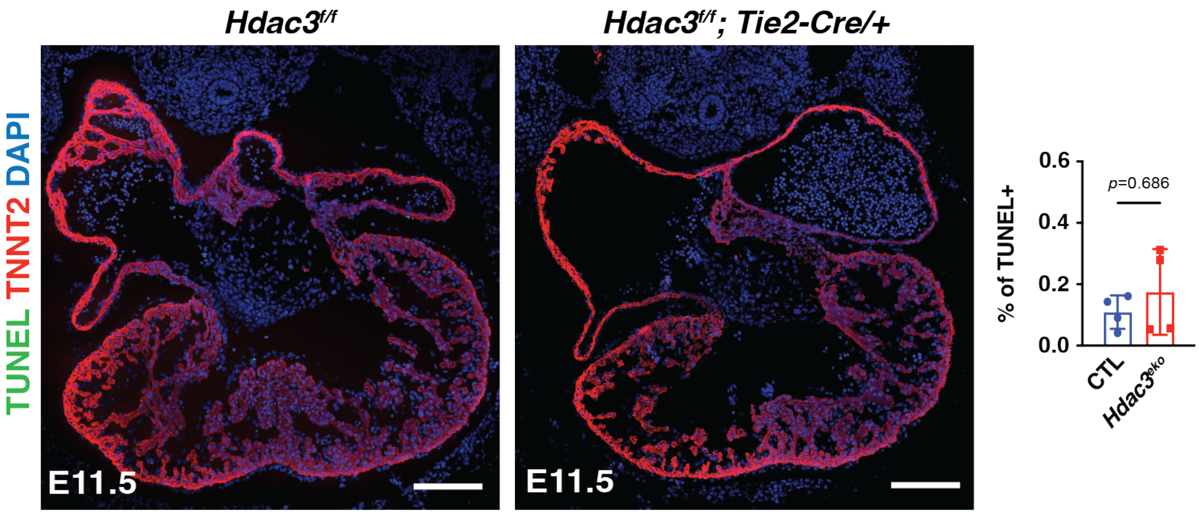


**Supplemental Figure 5. Apoptosis assessment in E11.5 *Hdac3*$^{tko}$ (*Hdac3*$^{f/f}$*; Tie2-Cre/+*) and littermate control (CTL, *Hdac3*$^{f/f}$) hearts by TUNEL staining.**

Scale bars, 100 µm. Quantitation is shown on the right (n=4 in each group). *P*-values were determined by the Mann-Whitney U test.

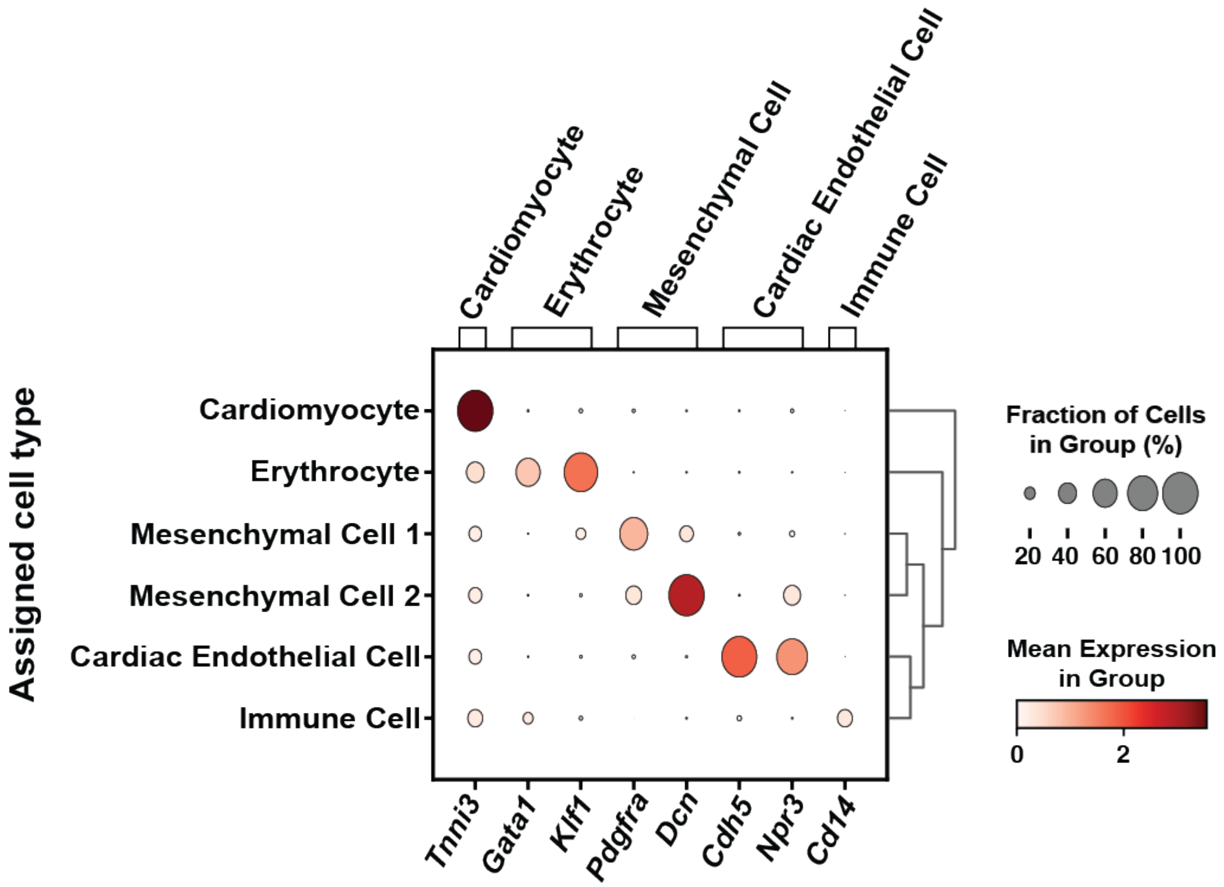


**Supplemental Figure 6. The cluster assignment of representative genes for various cardiac cell types.**

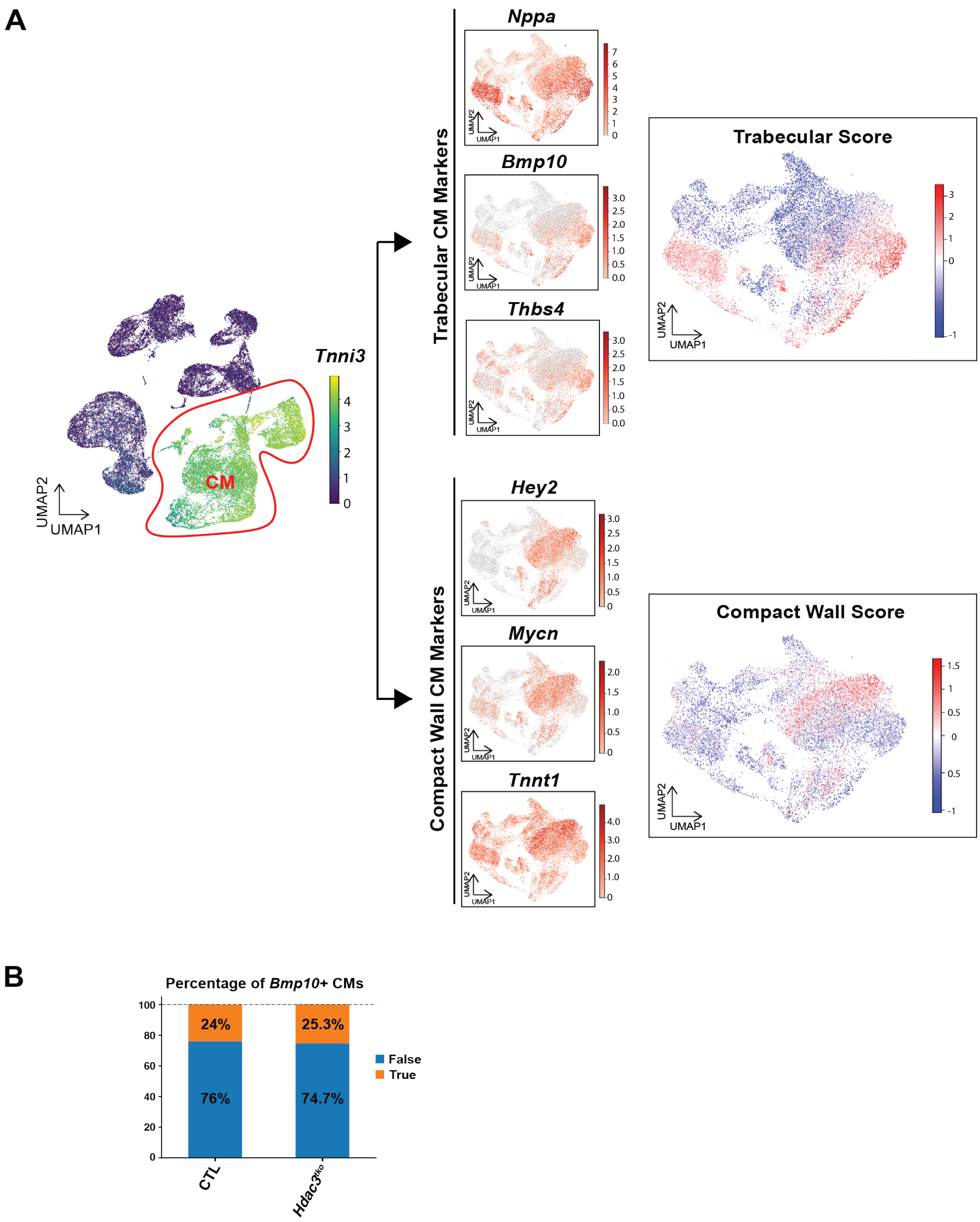


**Supplemental Figure 7. UMAP plots depicting the score of trabecular and compact wall cardiomyocytes and percentage of *Bmp10*+ Cells.**

**(A)** UMAP plots of trabecular score and compact wall score. Within *Tnni3*+ E11.5 cardiomyocytes (CMs), UMAP plots demonstrate the expression of trabecular CM marker genes including *Nppa*, *Bmp10* and *Thbs4*, and compact wall CM marker genes including *Hey2*, *Mycn* and *Tnnt1*. The calculated trabecular score and compact wall score are shown on the right. **(B)** Percentage of *Bmp10*+ cells within E11.5 *Hdac3*$^{tko}$ and littermate control (CTL) hearts. n=4 for each group.

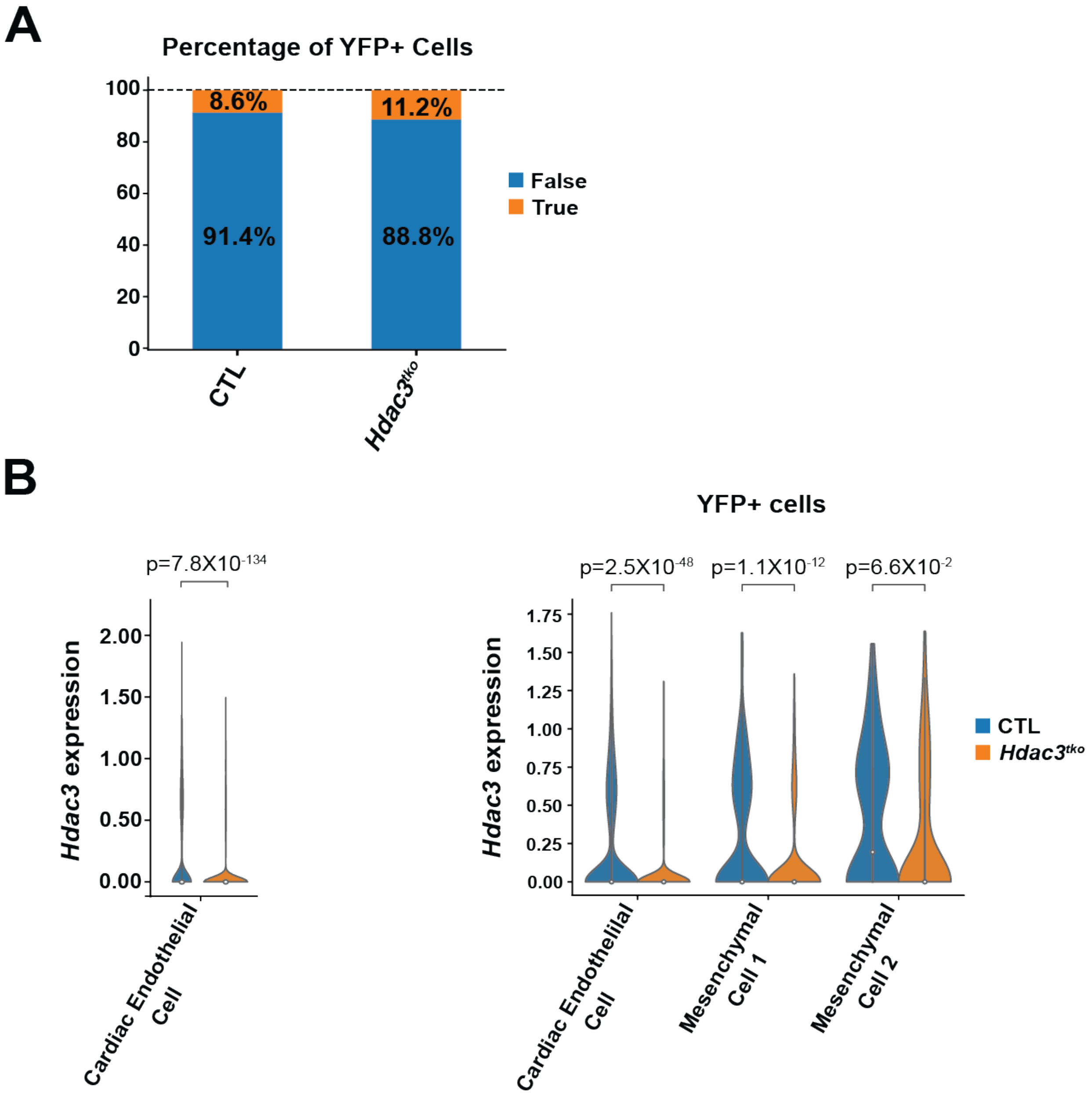


**Supplemental Figure 8. scRNA-seq analysis of YFP+ cells in E11.5 *Hdac3*$^{tko}$ and littermate control (CTL) hearts.**

**(A)** Percentage of YFP+ cells. CTL: n=3, *Hdac3*$^{tko}$: n=4. **(B)** Expression of *Hdac3* in the scRNA-seq data. (Left) Violin plots visualizing *Hdac3* gene expression in the endocardium. (Right) *Hdac3* expression in YFP+ clusters. *P*-value was determined by the Mann-Whitney U test.

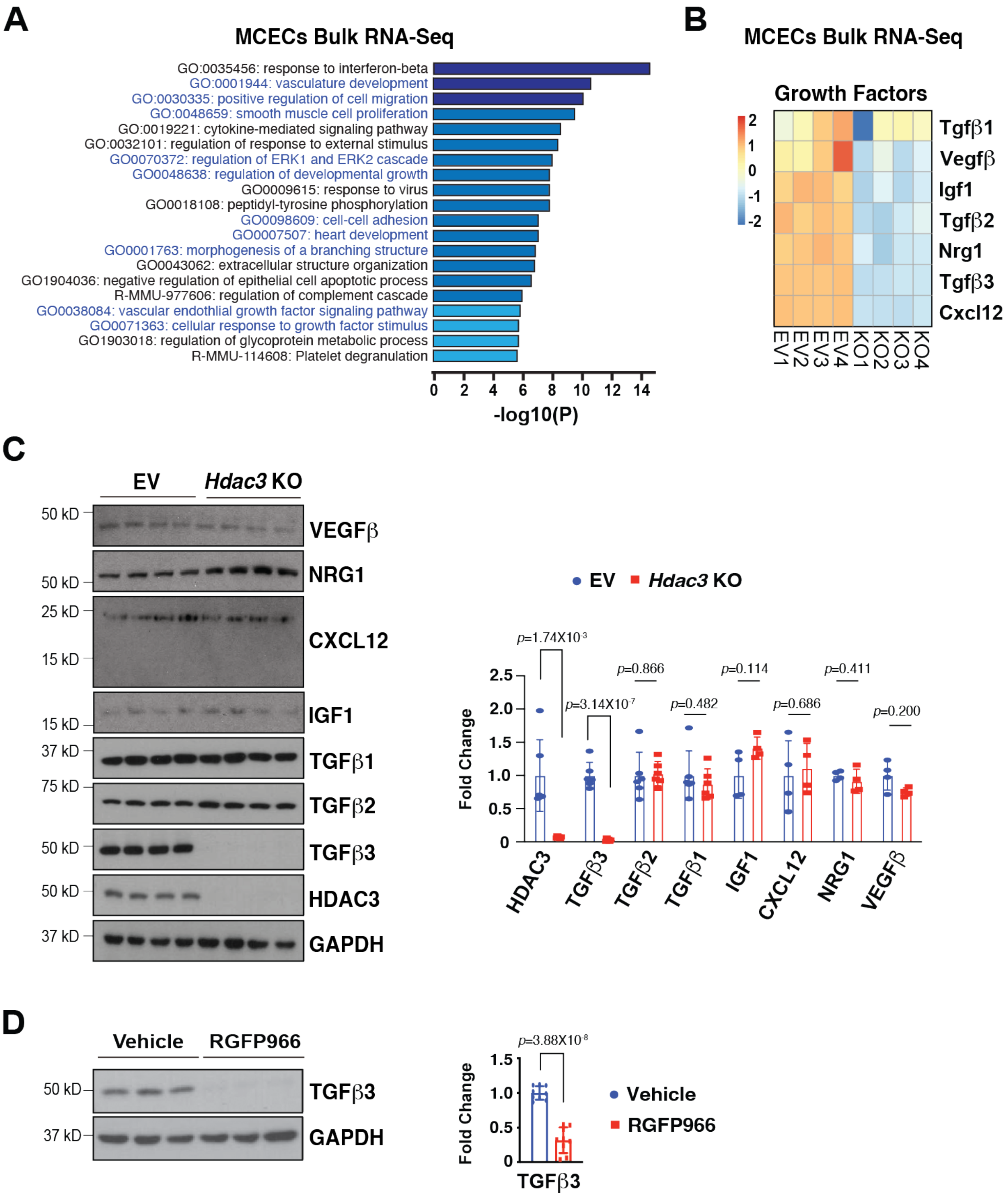


**Supplemental Figure 9. HDAC3 induces the expression of Tgfß3 dependent on its deacetylase activity.**

**(A)** Gene Ontology (GO) pathway analyses of RNA Seq in *Hdac3* KO and EV MCECs. n=4 in each group. Cut-off criteria: adjusted *P*-value<0.01. **(B)** Heatmap of growth factors (*Tgfβ1*, *Vegfβ*, *Igf1*, *Tgfβ2*, *Nrg1*, *Tgfβ3* and *Cxcl12*) in *Hdac3* KO and CTL MCECs. Data were extracted from the bulk RNA-sequencing data. **(C)** Quantification of TGFβ1, VEGFβ, IGF1, TGFβ2, NRG1, TGFβ3 and CXCL12 in *Hdac3* KO MCECs by western blot. GAPDH was used as protein loading control. n=6 for HDAC3, TGFβ1, TGFβ2 and TGFβ3 in each group. n=4 for VEGFβ, IGF1, NRG1 and CXCL12 in each group. *P*-values were determined by unpaired two tailed Student's *t*-test and the Mann-Whitney U test. **(D)** Reduced TGFβ3 protein expression after RGFP966 (selective HDAC3 inhibitor) treatment. MCEC were treated with 10 μM RGFP966 or vehicle for 24 hours. n=9 in each group. *P*-values were determined by unpaired two tailed Student's *t*-test.

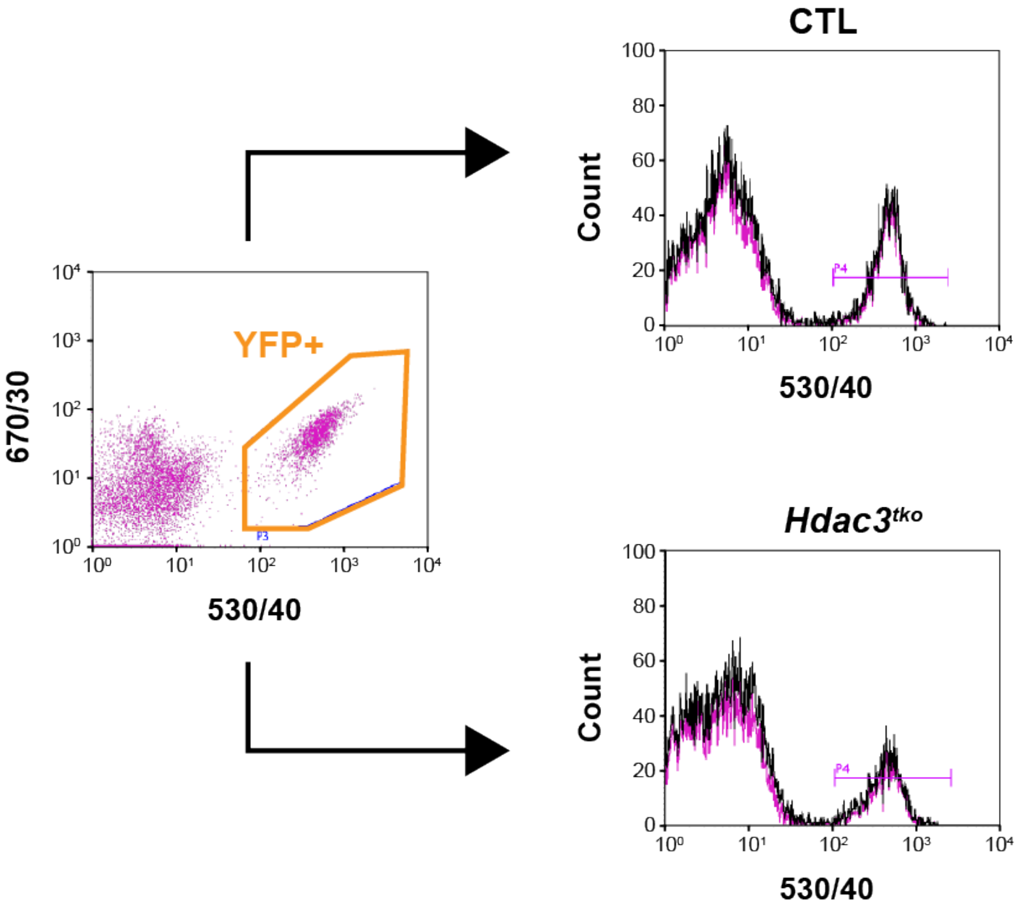


**Supplemental Figure 10. Fluorescence-activated cell sorting of YFP+ E11.5 cardiac endothelial cells.**

YFP+ cells from *Hdac3*$^{tko}$ (*Hdac3*$^{f/f}$*; Tie2-Cre/+; R26*$^{eYFP/+}$) and CTL (*Hdac3*$^{f/+}$*; Tie2-Cre/+; R26*$^{eYFP/+}$) hearts were gated and sorted for subsequent qRT-PCR analysis.

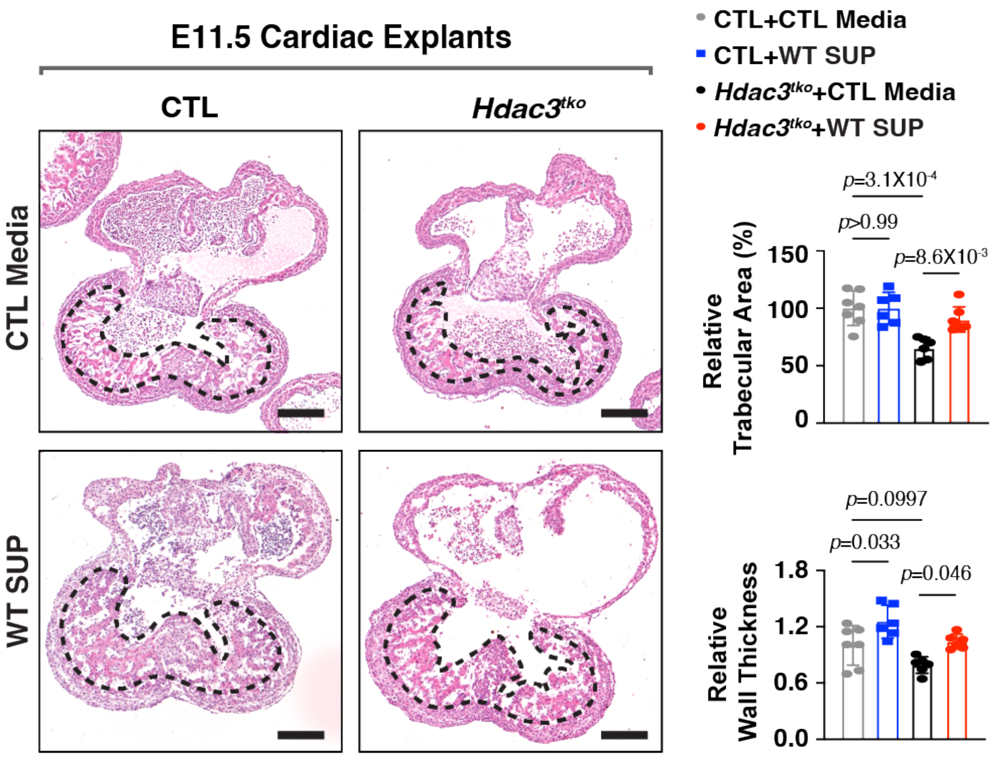


**Supplemental Figure 11. Wildtype MCEC supernatant supplementation rescues myocardial growth defects in endocardial *Hdac3* deficient hearts.**

Representative images of Hematoxylin and Eosin staining of E11.5 cardiac explants treated with wildtype (WT) MCEC medium (Final concentration of SUP: 250 ng/mL) or control (CTL) medium (DMEM/F-12 only) for 24 hours. Quantifications of relative ventricular trabecular area and wall thickness (relative to the mean values of CTL cardiac explants in CTL media group) are shown on the right. CTL ($Hdac3^{f/f}$) cardiac explants in CTL media: n=7, $Hdac3^{tko}$ ($Hdac3^{f/f}$; *Tie2-Cre/+*) cardiac explants in CTL media: n=6, CTL cardiac explants in WT MCEC supernatant (SUP) treatment: n=6, $Hdac3^{tko}$ cardiac explants in WT MCEC SUP treatment: n=6. Scale bars, 200 µm. *P*-values were determined by the one-way ANOVA followed by Tukey post hoc test.

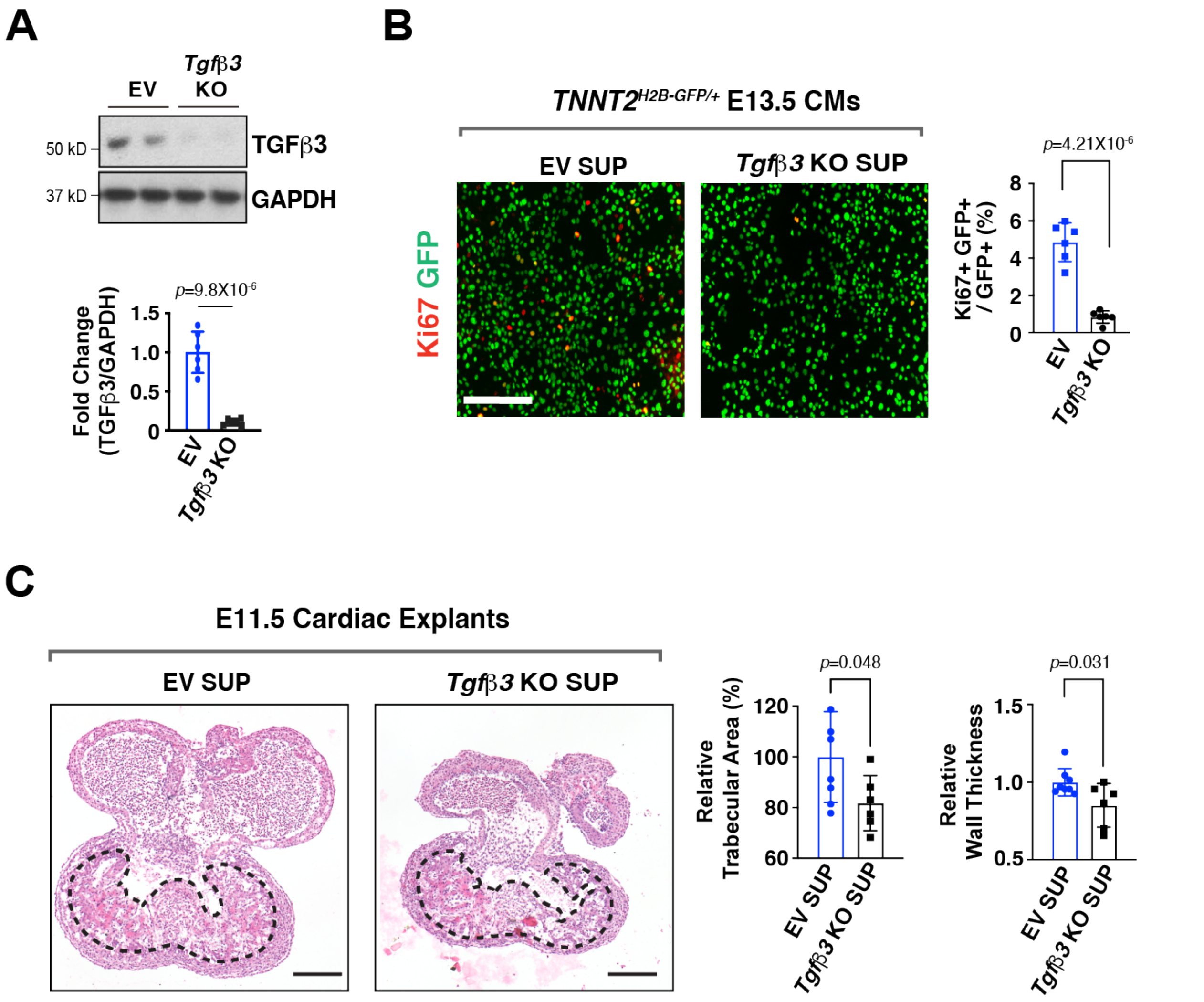


**Supplemental Figure 12. Reduced capacity of *Tgfβ3* knockout (KO) MCEC supernatants in inducing myocardial growth,**

**(A)** Generation of *Tgfβ3* KO and empty vector control (EV) MCECs by CRISPR/Cas9. Deletion of *Tgfβ3* was verified by western blot. Quantification is shown on the right (n=6 in each group). **(B)** The effects of *Tgfβ3* KO MCEC supernatants (SUPs) on E13.5 $Tnnt2^{nGFP/+}$ CM proliferation. Representative immunofluorescence micrographs are shown. Scale bar, 200 µm. Percentage of Ki67+ CMs were quantitated (n=6 in each group). **(C)** Representative images of Hematoxylin and Eosin staining of E11.5 wildtype heart explants (frontal section). Quantifications of relative ventricular trabecular area and wall thickness (relative to the mean values of the EV SUP group) are shown on the right. EV SUP: n=8, *Tgfβ3* KO MCEC SUP treatment: n=6. Scale bars, 200 μm. *P*-values were determined by unpaired two tailed Student's *t*-test.

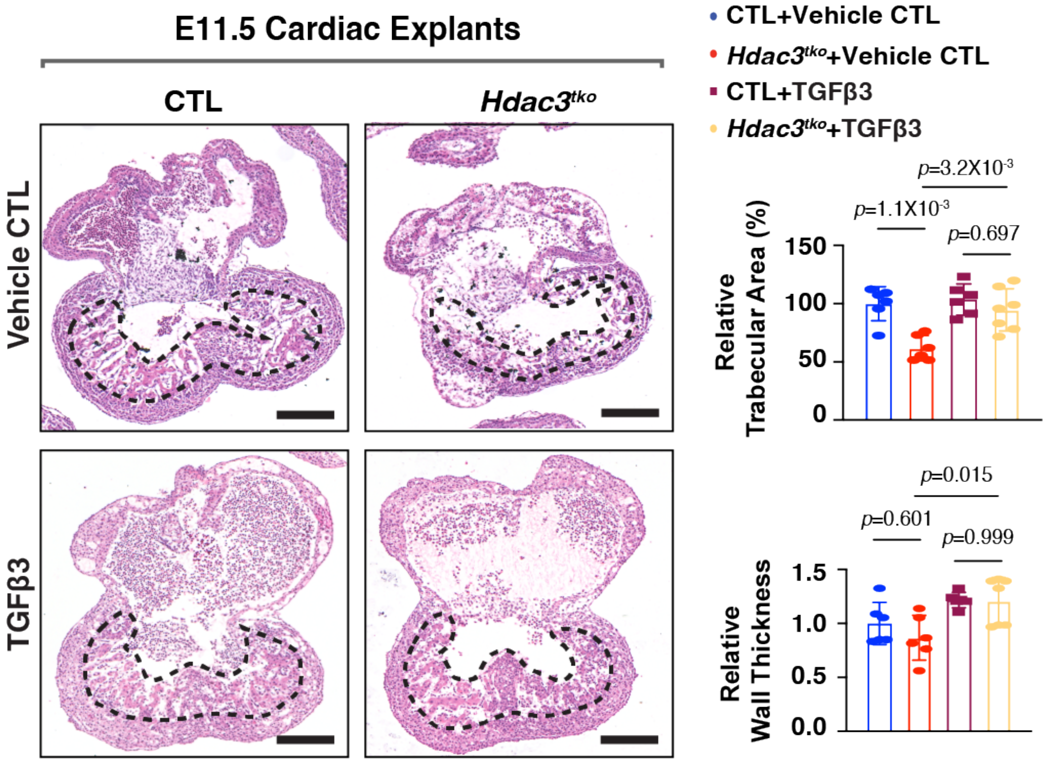


**Supplemental Figure 13. TGFβ3 supplementation rescues myocardial growth defects in endocardial *Hdac3* deficient hearts.**

Representative images of Hematoxylin and Eosin staining of E11.5 cardiac explants treated with TGFβ3 medium (TGFβ3 final concentration: 250 ng/mL) or vehicle control medium for 24 hours. Quantifications of relative ventricular trabecular area and wall thickness (relative to the mean values of CTL cardiac explants in Vehicle CTL media group) are shown on the right. CTL (*Hdac3*$^{f/f}$) cardiac explants in Vehicle CTL media: n=6, *Hdac3*$^{tko}$ (*Hdac3*$^{f/f}$*; Tie2-Cre/+*) cardiac explants in Vehicle CTL media: n=6, CTL cardiac explants in TGFβ3 media: n=6, *Hdac3*$^{tko}$ cardiac explants in TGFβ3 media: n=7. Scale bars, 200 µm. *P*-values were determined by the one-way ANOVA followed by Tukey post hoc test.

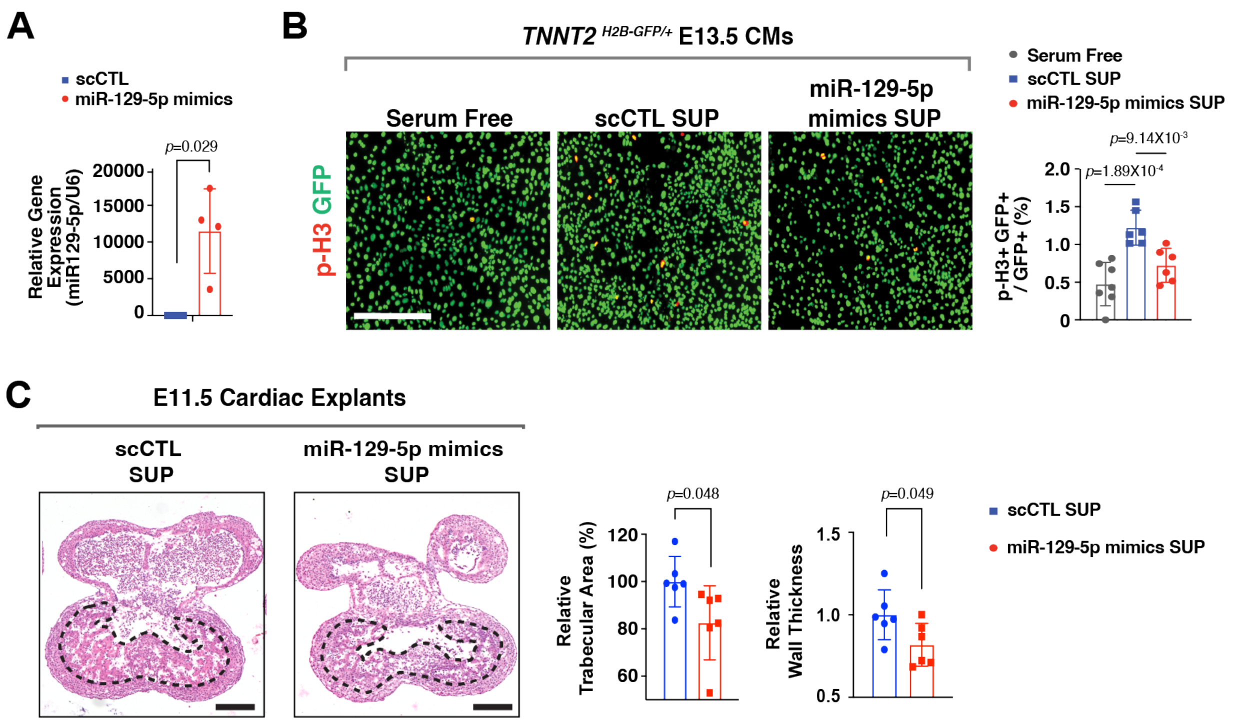


**Supplemental Figure 14. Reduced capacity of miR-129-5p-mimics-treated MCEC supernatants in inducing cardiomyocyte proliferation and myocardial growth.**

**(A)** Quantification of miR-129-5p expression in scramble control RNAs (scCTL) and miR129 mimics-treated MCECs. n=4 in each group. *P*-value was determined by the Mann-Whitney U test. **(B)** Evaluation of the effects of supernatants (SUPs) from miR-129-5p-mimics-treated MCECs on E13.5 *Tnnt2*$^{nGFP/+}$ CM proliferation. Representative immunofluorescence micrographs are presented. Scale bars, 200 µm. Percentage of phospho-histone H3 (p-H3)+ CMs was quantified. Independent samples: n=6 in each group. *P*-values were determined by one-way ANOVA followed by the Tukey post hoc test. **(C)** Representative images of Hematoxylin and Eosin staining of E11.5 wildtype heart explants (frontal section). Quantifications of relative ventricular trabecular area and wall thickness (relative to the mean values of the scCTL group) are shown on the right.

scCTL SUP: n=6, miR-129-5p mimics SUP: n=6. Scale bars, 200 μm. *P*-values were determined by unpaired two tailed Student's *t*-test.

**Supplemental Table 1. Genotype distribution of *Hdac3*$^{tko}$ embryos and offspring (♂: *Hdac3*$^{f/+}$*; Tie2-Cre* X ♀: *Hdac3*$^{f/f}$)**

| | **Others (*Hdac3*$^{f/f}$, *Hdac3*$^{f/+}$, *Tie2-Cre; Hdac3*$^{f/+}$) [expected: 75%]** | ***Hdac3*$^{tko}$ (*Tie2-Cre; Hdac3*$^{f/f}$) [expected: 25%]** | ***p*-value** |
|---|---|---|---|
| E9.5 [n (%)] | 140 (72.9%) | 52 (27.1%) | 0.505 |
| E10.5 [n (%)] | 168 (78.6%) | 46 (21.4%) | 0.2364 |
| E11.5 [n (%)] | 370 (78.6%) | 101 (21.4%) | 0.0747 |
| E12.5 [n (%)] | 91 (87.5%) | 13 (12.5%) | 0.0032* |
| E14.5 [n (%)] | 114 (99.1%) | 1 (0.9%) | <0.00001* |
| P0 | 336 (100%) | 0 | <0.00001* |

*P*-values were calculated by the comparison of genotype distribution and expected Mendelian Ratio.

**Supplemental Table 2. Genotype distribution of *Hdac3*$^{nko}$ embryos and offspring (♂: *Hdac3*$^{f/+}$; *Nfatc1*$^{Cre/+}$ X ♀: *Hdac3*$^{f/f}$)**

| | **Others (*Hdac3*$^{f/f}$, *Hdac3*$^{f/+}$, *Nfatc1*$^{Cre/+}$; *Hdac3*$^{f/+}$) [expected: 75%]** | ***Hdac3*$^{nko}$ (*Nfatc1*$^{Cre/+}$; *Hdac3*$^{f/f}$) [expected: 25%]** | ***p*-value** |
|---|---|---|---|
| E14.5 [n (%)] | 66 (82.5%) | 14 (17.5%) | 0.1213 |
| P0 | 209 (100%) | 0 | <0.00001* |

*P*-values were calculated by the comparison of genotype distribution and expected Mendelian Ratio.
